## Supplementary Figs for "Apoptosis-mediated ADAM10 activation removes a mucin barrier promoting T cell efferocytosis"

Linnea Drexhage *et al*

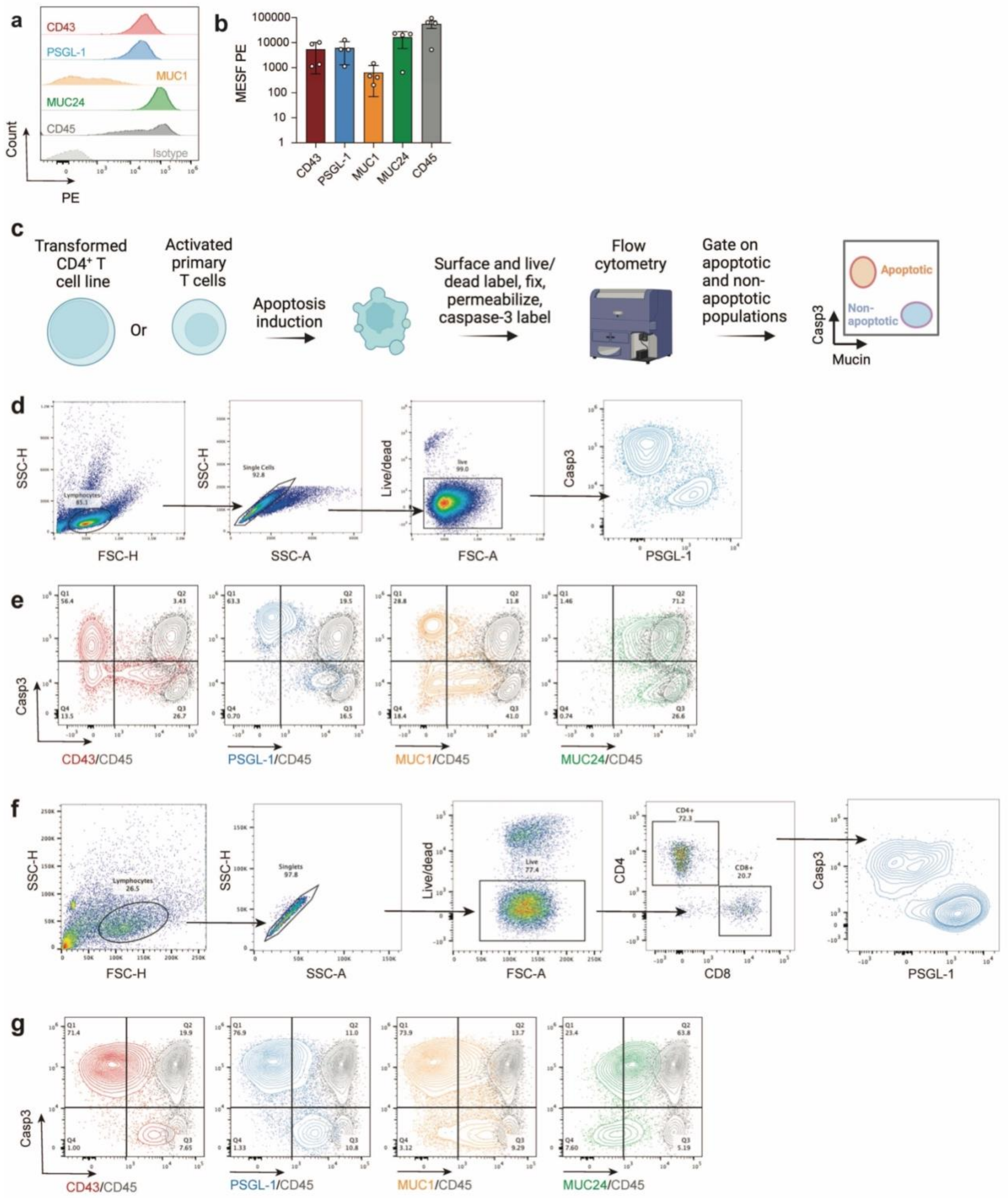


**Supplementary Fig. 1. Analysis of mucin expression on the surface of apoptotic T cells.** **a,** Relative cell surface mucin expression on CEM analyzed by flow cytometry. **b,** Number of mucin molecules estimated per CEM cell using Molecules of Equivalent Soluble Fluorochrome (MESF) quantification, $n$ = 4 independent experiments. **c,** Workflow for apoptosis induction, detection and mucin expression on healthy and apoptotic T cells. **d,** Flow cytometry gating strategy for analysis of mucin expression on apoptotic and non-apoptotic T cells; side-scatter = SSC, forward scatter = FSC. **e**, Dot plot of activated caspase-3 (Casp3) vs mucin expression on the surface of staurosporine-treated CEM, all mucins compared to CD45. **f**, Flow cytometry gating strategy for analysis of mucin expression on apoptotic and non-apoptotic primary T cells gated on either $CD4^+$ or $CD8^+$ populations. **g,** Dot plot of activated caspase-3 (Casp3) vs mucin expression on the surface of activated staurosporine-treated primary $CD4^+$ T cells, all mucins compared to CD45.

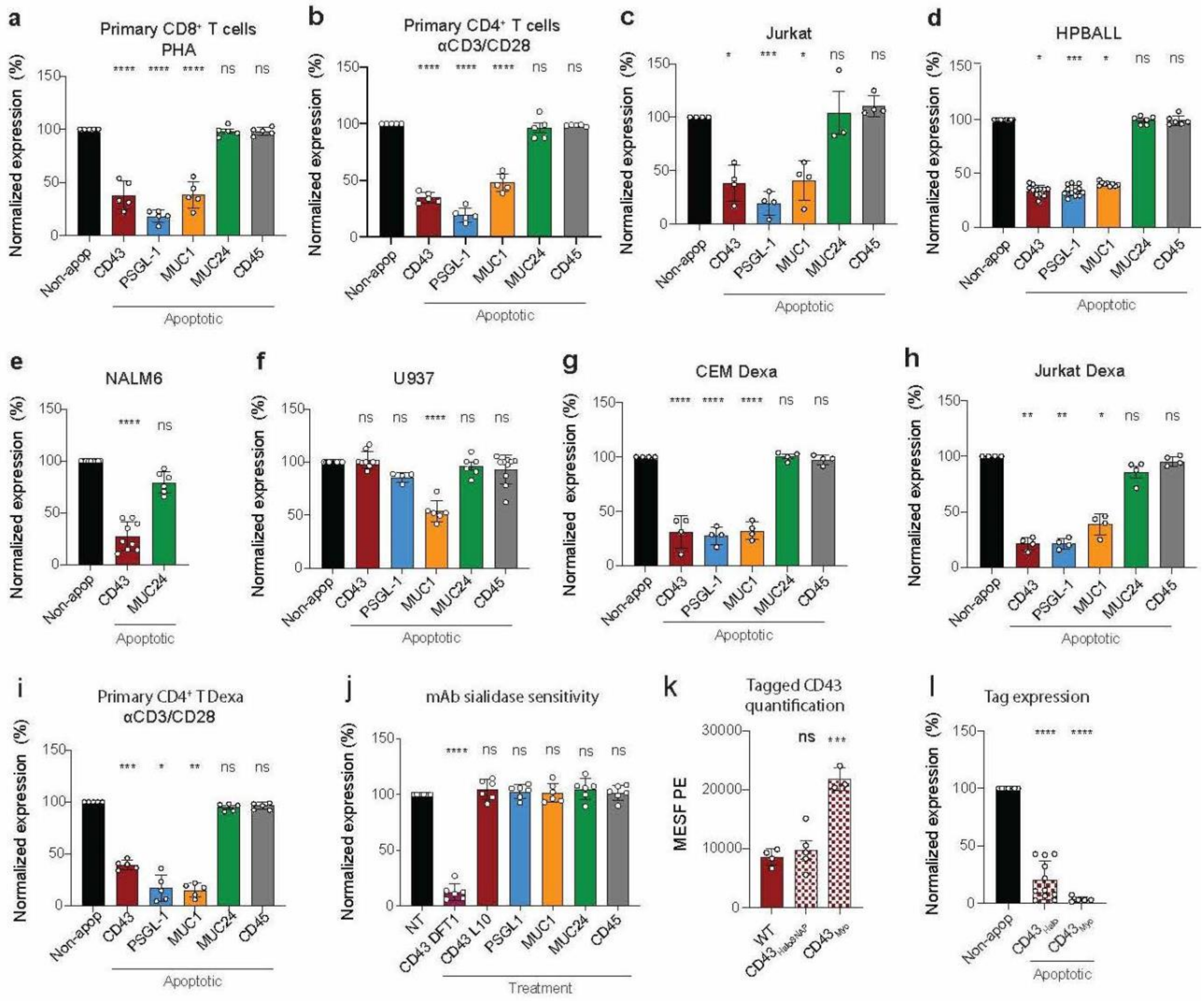


**Supplementary Fig. 2. Mucin expression on apoptotic immune cells**. **a**, Mucin expression on primary CD8$^+$ T cells activated with PHA+IL-2 and induced for apoptosis with staurosporine; n = 5 independent donors. **b**, Mucin expression on primary CD4$^+$ T cells activated with CD3/CD28 beads + IL-2 and induced for apoptosis with staurosporine; *n* = 5 independent donors. **c**, Mucin expression on Jurkat T cells induced for apoptosis with staurosporine; *n* = 4 independent experiments. **d**, mucin expression on HPBALL induced for apoptosis with staurosporine; *n* = 6 - 14 independent experiments. **e**, Mucin expression on NALM6 induced for apoptosis with staurosporine; *n* = 6 - 9 independent experiments. **f**, Mucin expression on U937 induced for apoptosis with staurosporine; *n* = 6 - 10 independent experiments. **g**, Mucin expression on CEM induced for apoptosis using dexamethasone (Dexa); *n* = 4 independent experiments. **h**, Mucin expression on Jurkat induced for apoptosis using Dexa; *n* = 4 independent experiments. **i**, Mucin expression on primary CD4$^+$ T cells activated with CD3/CD28 + IL-2 and induced for apoptosis with Dexa; *n* = 5 independent donors. **j**, Mucin-specific antibody binding to sialidase-treated CEM, with data normalized to mucin expression on non-sialidase-treated (NT) cells set at 100% and shown as a single bar; *n* = 3 independent experiments. **k**, Relative CD43$_{Myc}$ and CD43$_{Halo}$ expression levels determined by MESF on CEM-CD43$_{KO}$ cells stably transfected with an N-terminal Halo and C-terminal SNAP tag (CD43$_{Halo/SNAP}$) or N-terminal Myc tag (CD43$_{Myc}$); *n* = 3 - 5 independent experiments. **l**, Loss of CD43$_{Myc}$ and CD43$_{Halo}$ labels during apoptosis relative to non-apoptotic counterparts, both normalized to 100% shown as a single bar; *n* = 5 - 12 independent experiments. **p*<0.05; ***p*<0.01, ****p*<0.001, *****p*<0.0001; ns = not significant.

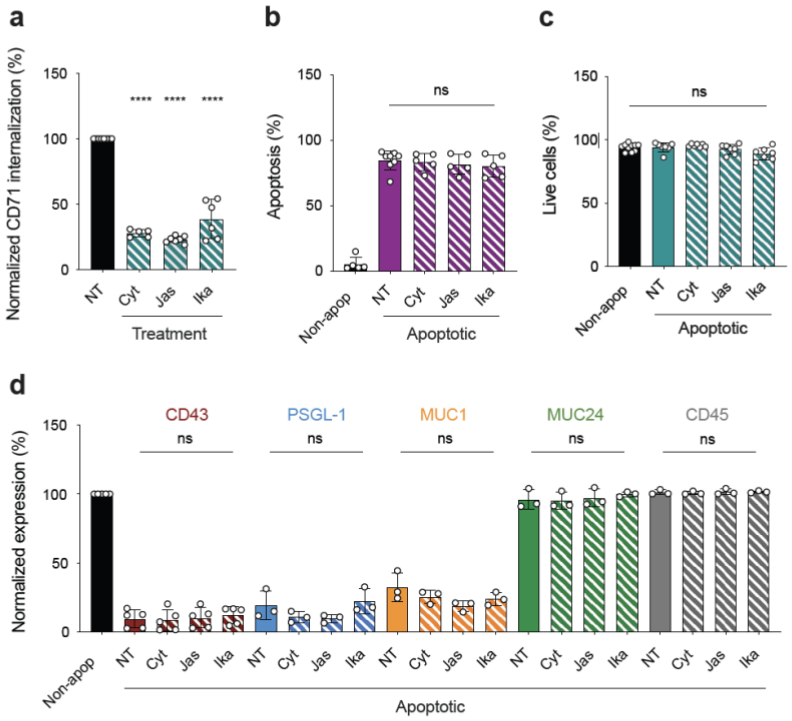


**Supplementary Fig. 3. Inhibition of actin remodeling and endocytosis does not prevent mucin loss. a,** Effect of actin remodeling inhibitors cytochalasin-D (Cyt), Jasplakinolide (Jas) and endocytosis inhibitor Ikarugamycin (Ika) on antibody-mediated CD71 internalization, non-treated (NT) cells normalized to 100%; $n$ = 5 - 8 independent experiments. **b,** Effect of inhibitors on staurospaurine-induced apoptosis assayed by activated caspase-3 labeling, $n$ = 5 - 8 independent experiments. **c,** Lack of inhibitor-mediated cytotoxicity, live (apoptotic and non-apoptotic) cells gated using live/dead stain, $n$ = 5 - 7 independent experiments. **d,** Lack of effect of inhibitors on mucin loss during apoptosis, $n$ = 3 - 5 independent experiments. **** $p$<0.0001, ns = non-significant.

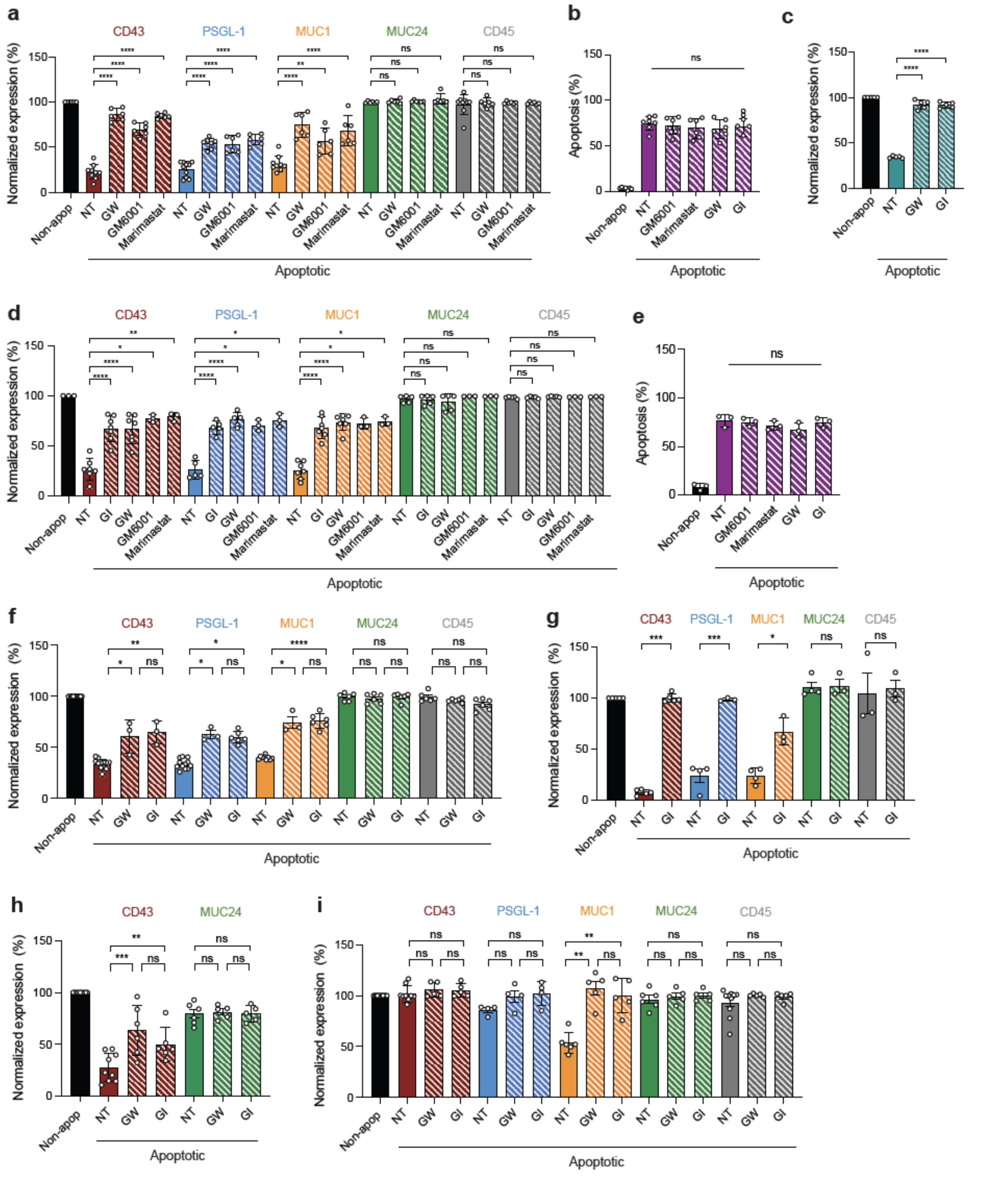


**Supplementary Fig. 4. Metalloprotease inhibitors reduce mucin shedding. a**, Metalloprotease inhibitors were added to staurospaurine-treated CEM and cells labelled for activated caspase-3 and mucin expression, non-apoptotic cells (Non-apop) normalized to 100% shown as a single bar, NT = non-treated; *n* = 6 - 11 independent experiments. **b**, Effect of metalloprotease inhibitors on staurosporine-induced CEM apoptosis; *n* = 6 - 7 independent experiments. **c,** CD46 expression on staurosporine-treated CEM, pre-treated or NT with metalloprotease inhibitors, non-apoptotic (Non-apop) cells normalized to 100% shown as a single bar; *n* = 5 independent experiments. **d**, Mucin expression on staurospaurine-treated primary CD4$^{+}$ T cells pre-treated or untreated (NT) with metalloprotease inhibitors, non-apoptotic (Non-apop) cells normalized to 100% shown as a single bar; *n* = 3 - 9 independent donors. **e**, Effect of metalloprotease inhibitors on staurosporine-induced primary T cell apoptosis determined by activated caspase-3 labeling; *n* = 3 independent experiments. **f**, Mucin expression on staurosporine-treated HPBALL, pre-treated or untreated (NT) with metalloprotease inhibitors GW or GI, non-apoptotic (Non-apop) cells normalized to 100% shown as a single bar; *n* = 3 - 12 independent experiments. **g**, Mucin expression on staurosporine-treated Jurkat, pre-treated or NT with ADAM10 inhibitor GI, non-apoptotic (Non-apop) cells normalized to 100% shown as a single bar; *n* = 3-5 independent experiments. **h**, Mucin expression on staurosporine-treated NALM6, pre-treated or not treated (NT) with metalloprotease inhibitors GW or GI, non-apoptotic (Non-apop) cells normalized to 100% shown as a single bar; *n* = 5 - 9 independent experiments. **i**, Mucin expression on staurosporine-treated U937, pre-treated or untreated (NT) with ADAM10 inhibitor GI, non-apoptotic (Non-apop) cells normalized to 100% shown as a single bar; *n* = 4 - 8 independent experiments. *$p$<0.05; **$p$<0.01, ***$p$<0.001; ****$p$<0.0001; ns = not significant.

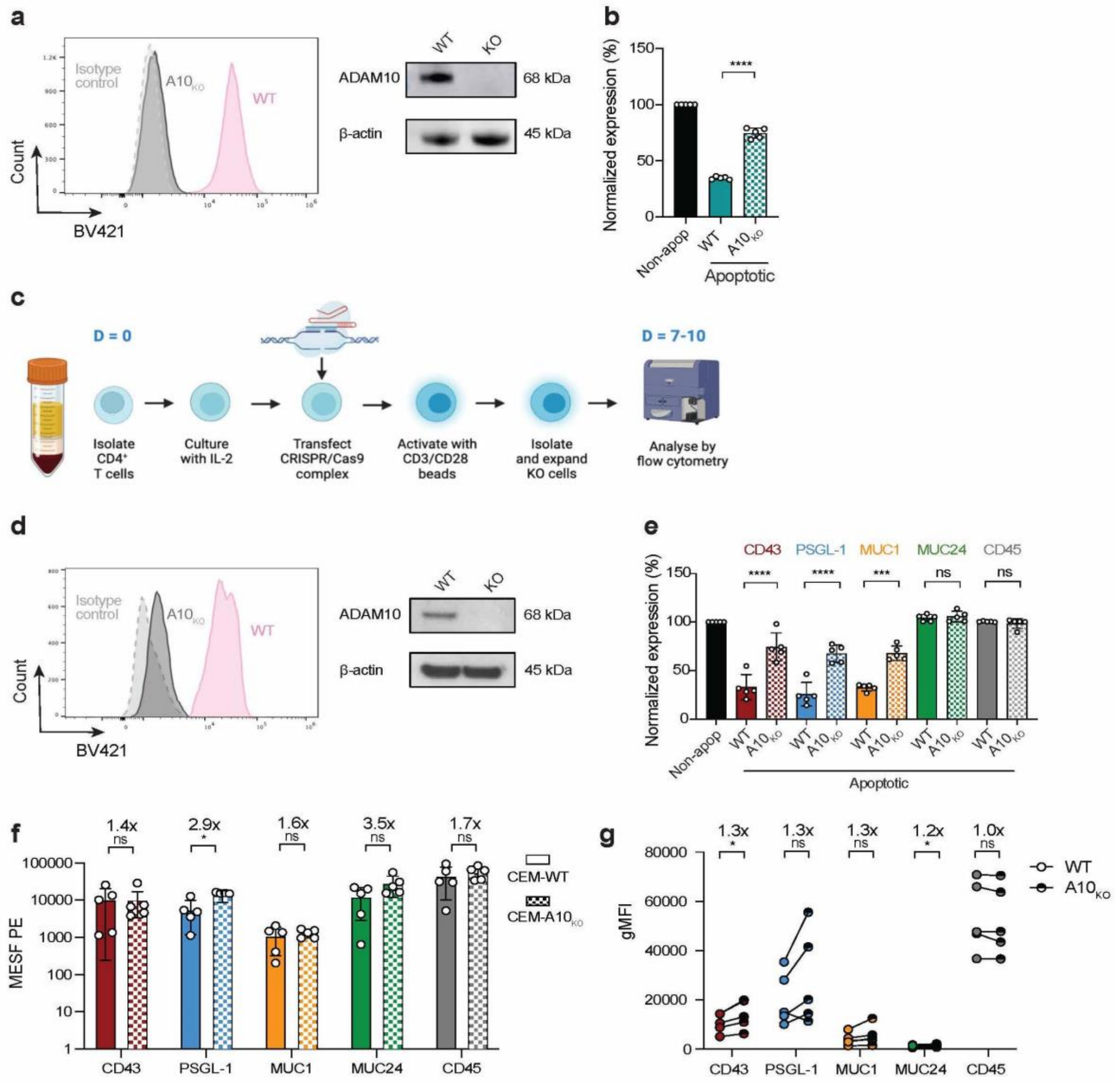


**Supplementary Fig. 5. CRISPR-Cas9 knock out of ADAM10 inhibits mucin shedding. a,** Analysis of cell surface ADAM10 expression, CEM WT and CRISPR-Cas9 KO line $ADAM10_{KO}$ = $A10_{KO}$, dotted grey line = isotype control, and western blot of CEM lysate for $A10_{KO}$ compared to WT CEM, β-actin = loading control, molecular weight (kDa) of band to right of blot. **b,** Cleavage of CD46 in apoptotic CEM WT and $A10_{KO}$, non-apoptotic cells (Non-apop) normalized to 100% shown as a single bar, $n$ = 3 independent experiments. **c,** Workflow for ADAM10 CRISPR-Cas9 KO in primary $CD4^+$ T cells. **d,** Flow cytometric cell surface and western blot analysis of CRISPR-Cas9 $A10_{KO}$ primary $CD4^+$ T cells treated with staurosporine and analyzed by flow cytometry, and western blot of T cell lysate for $A10_{KO}$ compared to WT T cells, β-actin = loading control, molecular weight (kDa) of band to right of blot. **e**, Staurosporine-treated WT and $A10_{KO}$ primary T cells analyzed for mucin expression, non-apoptotic cells (Non-apop) normalized to 100% and represented as a single bar for all groups; $n$ = 5 independent experiments. **f**, MESF analysis of mucin expression on healthy CEM WT and $A10_{KO}$, numbers above bars reflect fold-increase in expression; $n$ = 5 independent experiments. **g**, Flow cytometric analysis of mucin expression on healthy primary $CD4^+$ T cells expressed as geometric mean fluorescence (gMFI), numbers above bars reflect fold-increase in expression; $n$ = 5 independent donors. *$p<0.05$; ****$p<0.0001$; ns = not significant.

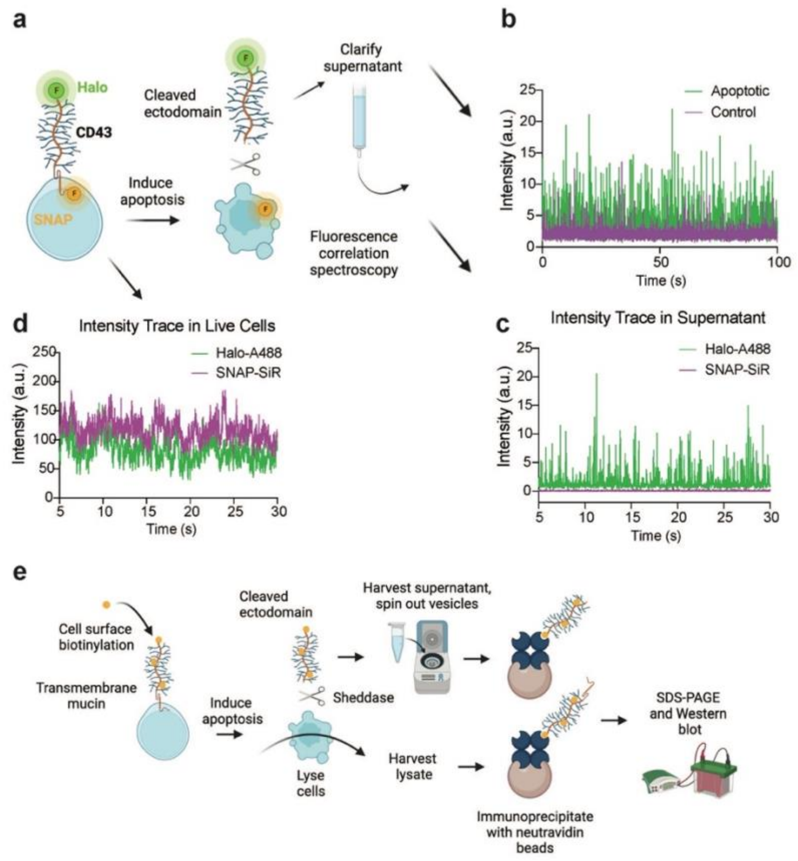


**Supplementary Fig. 6. Mucin ectodomains are shed from apoptotic cells.** **a,** Workflow for treatment of CEM$_{Halo/SNAP}$ cells and Fluorescence Correlation Spectroscopy (FCS) analysis. **b,** Time-resolved fluorescent single-molecule Halo-A488 peaks observed in processed supernatant from apoptotic compared to non-apoptotic CEM. **c,** Time-resolved intensity trace in supernatant of apoptotic CEM of Halo-A488 and SNAP-SIR labels. **d,** Time-resolved intensity trace of Halo-A488 and SNAP-SIR fluorescence in healthy CEM$_{Halo/SNAP}$ cells. **e,** Workflow for cell surface biotinylation and western blot analysis of cell-associated and soluble ectodomain mucin expression.

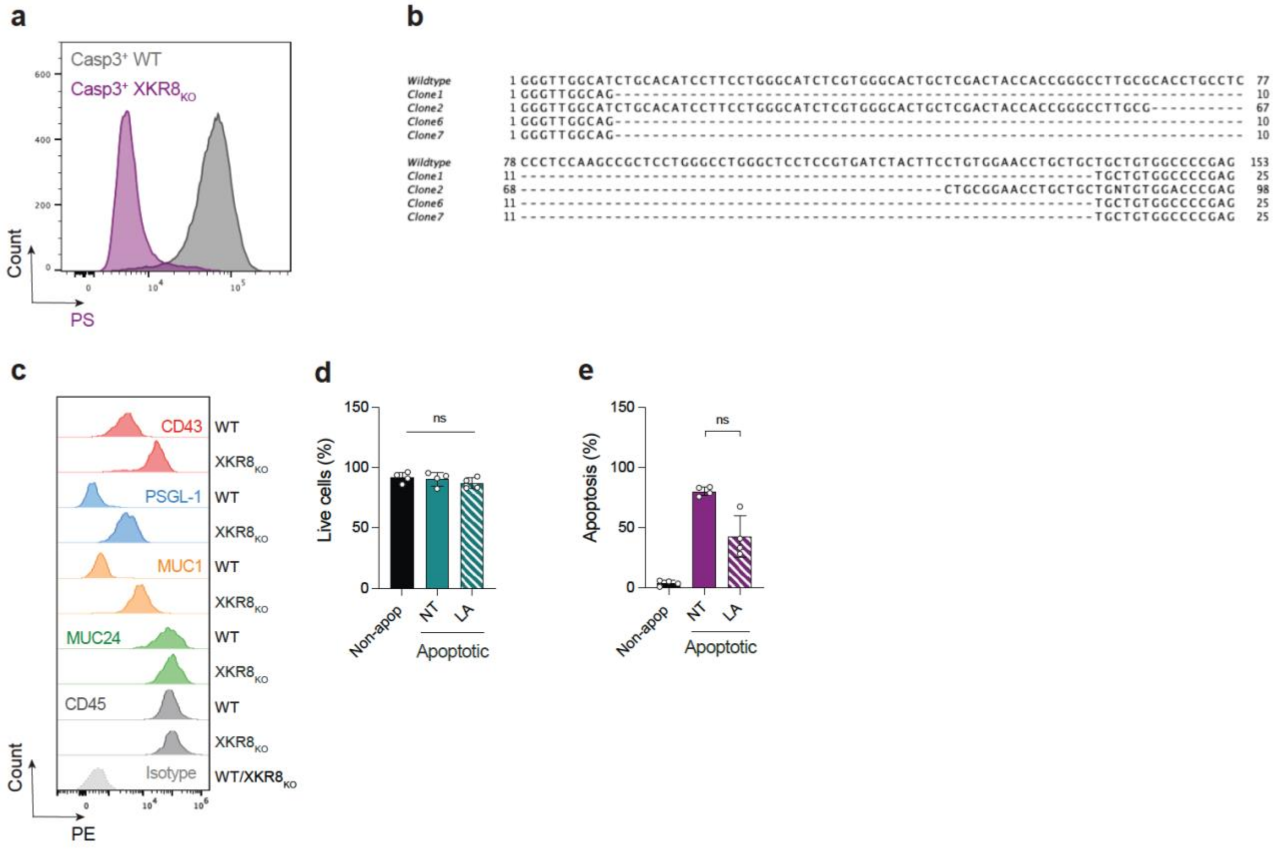


**Supplementary Fig. 7. XKR8$_{KO}$ and inhibition of PS-ADAM10 interaction. a,** Apoptosis induction in XKR8-KO (XKR8$_{KO}$) CEM clone does not flip PS assayed by annexin-V labelling in caspase-3-positive apoptotic cells. **b,** Sequence of XKR8$_{KO}$ PCR products to confirm deletions in Exon 3 of 4 knockout clones compared to WT CEM. **c,** Flow cytometric analysis showing XKR8$_{KO}$ in apoptotic CEM inhibits loss of CD43, PSGL-1 and MUC1. **d,** LA is non-cytotoxic to apoptotic CEM cells; *n* = 4 independent experiments. **e,** LA does not interfere with staurosporine-induced apoptosis in CEM cells, apoptosis quantified by detection of activated caspase-3 in fixed, permeabilized cells. ns = not significant; *n* = 4 independent experiments.

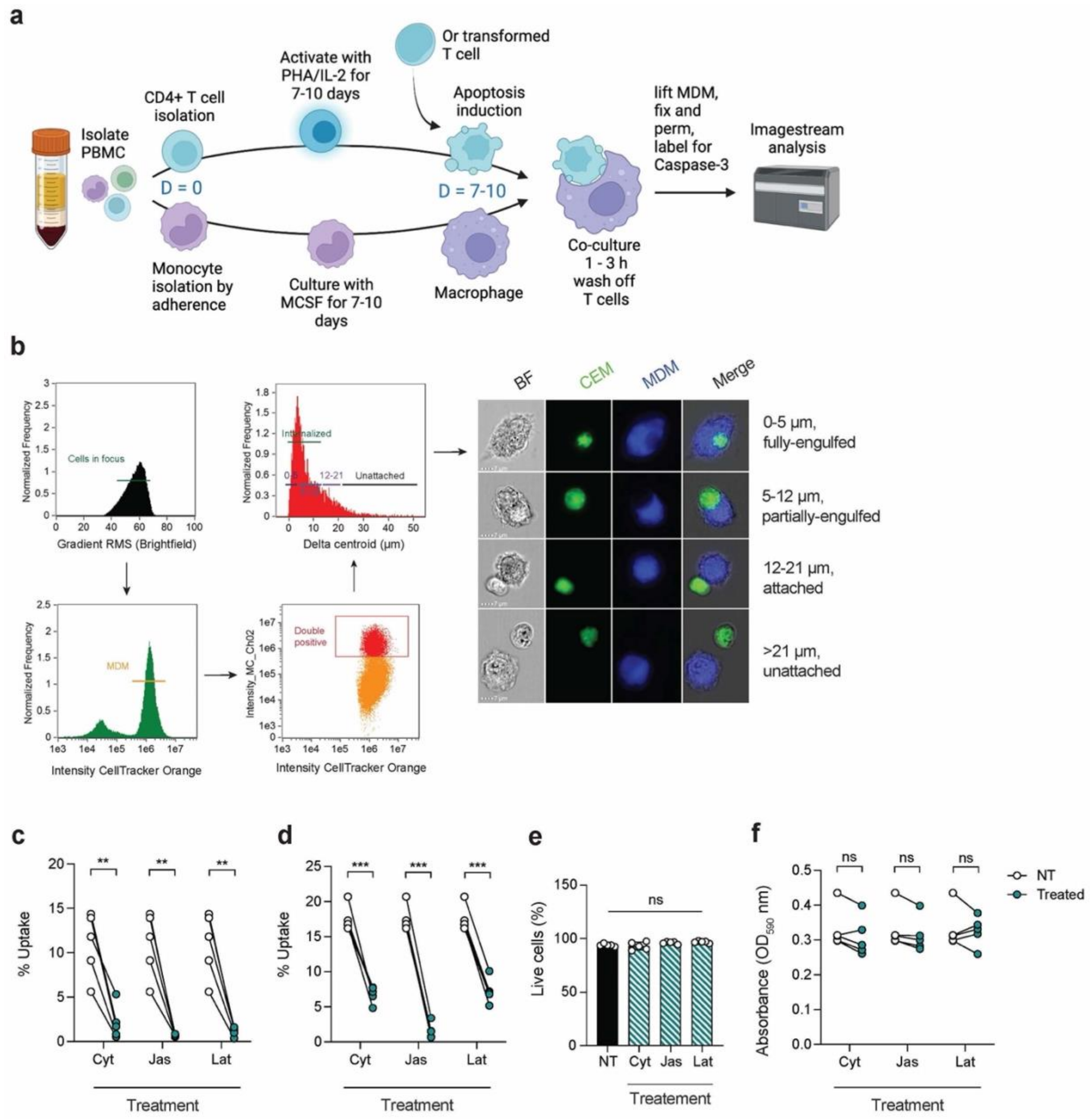


**Supplementary Fig. 8. Imagestream-based efferocytosis assay. a,** Processing workflow for peripheral blood mononuclear cells (PBMC) to obtain activated CD4$^{+}$ T cells and monocyte-derived macrophages (MDM), coculture and Imagestream analysis of T cell uptake. **b**, Imagestream gating strategy comprising selection of in-focus cells, selection of MDM, gating on double-positives and incorporation of the delta-centroid function with example images, bright-field = BF. Scale bar = 7 µm, distances are in µm. **c**, Pre-treatment (green circles) or not (open circles) of MDM with actin remodelling inhibitors cytochalasin-D (Cyt), jasplakinolide (Jas) or latrunculin (Lat) prior to coculture with apoptotic CEM, and Imagestream analysis for T cell uptake; $n$ = 5 independent experiments. **d**, Pre-treatment or not of MDM with actin remodelling inhibitors prior to coculture with fluorescent beads and flow cytometry analysis for bead uptake; $n$ = 5 independent experiments. **e**, Flow cytometric analysis of live/dead viability labelling of untreated (NT) and actin remodelling inhibitor-treated MDM. **f**, MTT assay for metabolic function of untreated (NT) and actin remodelling inhibitor-treated MDM. **$p$<0.01; ***$p$<0.001; ns = not significant.

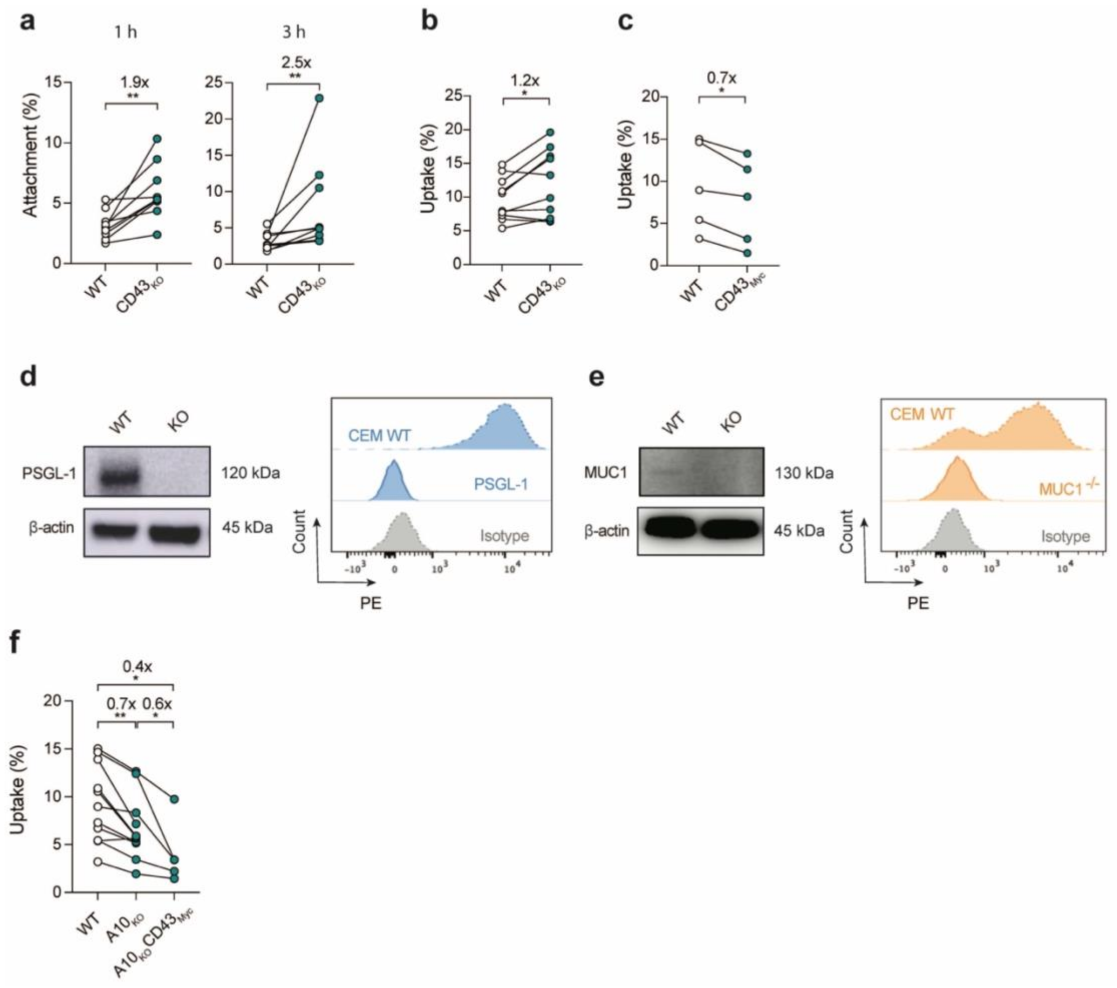


**Supplementary Fig. 9. Mucin loss enhances attachment to, and uptake of, apoptotic T cells.** **a,** Imagestream quantification of apoptotic WT (open circles) and $CD43_{KO}$ (green circles) CEM attachment to MDM at T = 1 and 3 h; $n$ = 6 - 9 independent experiments. **b,** Imagestream quantification of apoptotic CEM cell uptake by MDM after 3 h coculture comparing WT (open circles) and $CD43_{KO}$ (green circles); $n$ = 10 independent experiments. **c**, Imagestream quantification of apoptotic CEM cell uptake by MDM after 3 h coculture comparing WT (open circles) and $CD43_{KO}$ overexpressing Myc-tagged CD43 ($CD43_{Myc}$, green circles). **d**, CRISPR-Cas9 KO of PSGL-1, western blot with β-actin loading control with molecular weight (kDa) of band to right of blot, and cell surface flow cytometric analysis. **e,** CRISPR-Cas9 KO of MUC1, western blot with β-actin loading control with molecular weight (kDa) of band to right of blot, and cell surface flow cytometric analysis. **f,** Imagestream analysis of apoptotic CEM cell uptake by MDM after 3 h coculture comparing WT, $ADAM10_{KO}$ ($A10_{KO}$) and $A10_{KO}$ overexpressing Myc-tagged CD43 ($A10_{KO}CD43_{Myc}$); $n$ = 5 - 10 independent experiments. *$p$<0.05; **$p$<0.01.

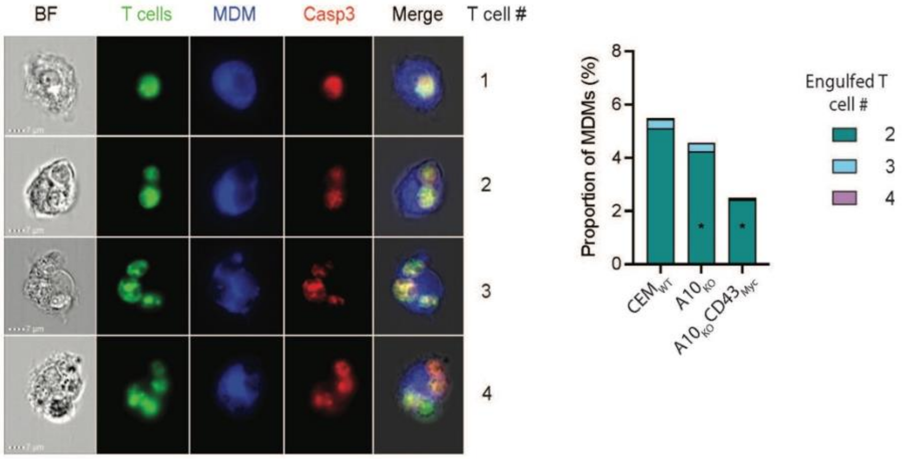


**Supplementary Fig. 10. MDM engulf multiple apoptotic CEM.** Staurosporine-treated CEM cells were washed, cocultured with MDM for 1 hr, washed in cold PBS/EDTA, lifted, fixed, permeabilized and labelled for CD3 (T cells) and caspase-3 (Casp3). Numbers of T cells in 1000 independent images double-positive for MDM and T cells were manually counted. MDM containing single T cells were excluded from the quantification to prioritize display of multiple uptake events. WT CEM ($CEM_{WT}$) were compared with CEM $A10_{KO}$ and CEM $A10_{KO}CD43_{Myc}$. *$p$<0.05.

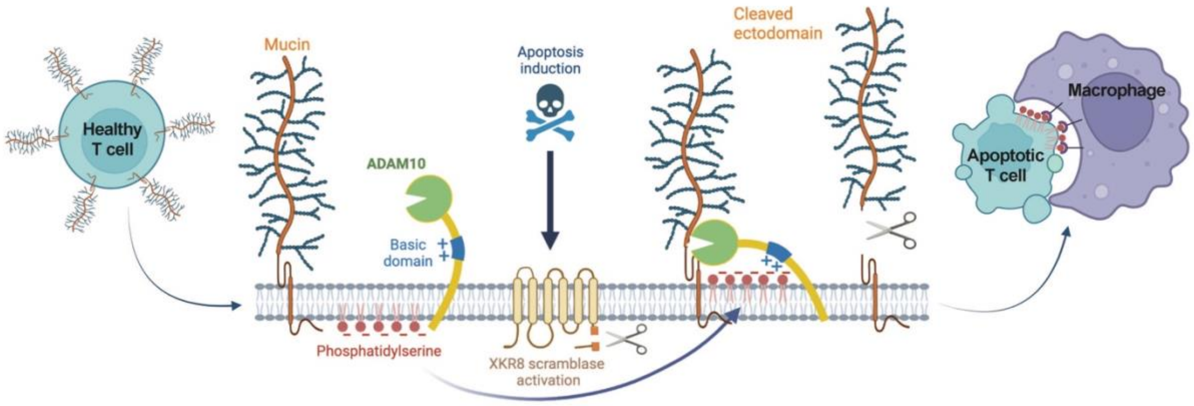


**Supplementary Fig. S11. Model for mucin loss facilitating T cell efferocytosis by macrophages.** Healthy T cells express a high density of extended transmembrane mucins in their glycocalyx. Activation of apoptosis leads to flipping of phosphatidylserine (PS) via caspase-3 activation of the scramblase XKR8, which in turn activates ADAM10 proteolytic activity for selected mucins. Mucin cleavage exposes membrane-proximal eat-me signals such as PS, which are recognised by cognate phagocytic receptors leading to efficient efferocytosis.
